# Fast and Accurate Exhaustive Higher-Order Epistasis Search with BitEpi

**DOI:** 10.1101/858282

**Authors:** Arash Bayat, Brendan Hosking, Yatish Jain, Cameron Hosking, Milindi Kodikara, Daniel Reti, Natalie A. Twine, Denis C. Bauer

**Affiliations:** Transformations Bioinformatics, Health and Biosecurity, Commonwealth Scientific and Industrial Research Organisation (CSIRO), North Ryde, New South Wales 2113, Australia; The Kinghorn Cancer Centre, Darlinghurst, New South Wales 2010, Australia; Department of Biomedical Sciences, Macquarie University, Macquarie Park, New South Wales 2113, Australia; Applied BioSciences, Faculty of Science and Engineering, Macquarie University, Macquarie Park, New South Wales 2113, Australia

## Abstract

**Motivation:** Complex genetic diseases may be modulated by a large number of epistatic interactions affecting a polygenic phenotype. Identifying these interactions is difficult due to computational complexity, especially in the case of higher-order interactions where more than two genomic variants are involved.

**Results:** In this paper, we present BitEpi, a fast and accurate method to test all possible combinations of up to four bi-allelic variants (i.e. Single Nucleotide Variant or SNV for short). BitEpi introduces a novel bitwise algorithm that is 2.1 and 56 times faster for 3-SNV and 4-SNV search, than established software. The novel entropy statistic used in BitEpi is 44% more accurate to identify interactive SNVs, incorporating a *p*-value-based significance testing. We demonstrate BitEpi on real world data of 4,900 samples and 87,000 SNPs. We also present EpiExplorer to visualize the potentially large number of individual and interacting SNVs in an interactive Cytoscape graph. EpiExplorer uses various visual elements to facilitate the discovery of true biological events in a complex polygenic environment.

## 1 Introduction

Complex diseases often have a multi-genic component where the individual SNVs can both independently and interactively contribute to the disease [1]. The interactive effects are referred to as epistasis [2, 3]. Epistatic interactions involving three or more SNVs (higher-order) have been suggested to contribute to the ‘missing heritability’ problem in complex diseases [1, 2]. However, detecting such interactions is computationally challenging due to the exponential complexity of the problem [4, 5, 6, 7, 8]. Given a dataset with *n* SNVs, the exhaustive epistasis search with the order of *m* (number of interactive SNVs) requires 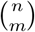 combinations of SNVs to be tested, resulting in a complexity of *O*(*n^m^*). For example, in a dataset with only 1,000 SNVs there are about 0.5, 166 and 41,417 million 2-SNV, 3-SNV and 4-SNV combinations to be tested respectively.

Due to the exponential complexity of higher-order exhaustive search algorithms, it is not practical to apply them to large datasets. However, it is possible to use a filter to reduce the search space to a smaller number of SNVs before a more in-depth analysis [9, 10, 11, 12]. Random Forest [13] is an efficient method for this filter as it preserves higher-order interactions [14]. Particularly, a new cloud-based implementation of Random Forest called VariantSpark [15] is able to process whole-genome data with 100,000,000 SNVs. It is capable of fitting tens of thousands of trees, which enables the interrogation of the search space more deeply, thereby reducing the chance of missing important interactions.

Irrespective of the applied filtering methodology, the key to discover and annotate a complete set of interactions is a fast exhaustive search. Non-exhaustive methods suffer from inaccuracy especially in case of “Strict and Pure” Higher-Order interactions [16] where none of the SNVs or subset of SNVs shows any association power. The association can be only detected when all interactive SNVs are considered together. There are several algorithms for finding pairwise (2-SNV) interactions between SNVs using exhaustive search approaches.

With execution time a major limitation, algorithmic improvements predominantly focus on speedup. For example, TEAM [17] uses a minimum spanning tree algorithm to minimize execution time. More recently, BOOST [18] delivered a 168-fold speed up over TEAM [5] by using bitwise operations for pairwise interactions.

However, as it is likely that more than two SNVs interact, efforts have been made to extend the exhaustive search capability to higher-order interactions. For example, CINOEDV [19] offers exhaustive searching for up to 5-SNV epistasis. However, with a focus on the visualization of the interactions, CINOEDV was not designed for speed and its non-parallel implementation in R is 66.5 times slower than BOOST when processing 100 SNVs [19] (for the 2-SNV search). Also the visualisation offered by CINOEDV is static and incapable of representing large interaction graphs. Capable of processing higher-order interactions more efficiently, MDR [20] (Multi-factor Dimensionality Reduction) is an extensive epistasis analysis platform offering parallel exhaustive search functionality. Improving on the algorithmic implementation further, MPI3SNP [21] adapts the bitwise approach used by BOOST. However, with MPI3SNP being limited to 3-SNV searches, the need for a fast higher-order search remains unaddressed.

In this paper, we introduce BitEpi, a fast and accurate exhaustive higher-order epistasis search program written in C++, which is able to test up to 4-SNV combination. BitEpi introduces a novel bitwise approach capable of handling higher-order interactions, making it the first bitwise optimization method to be able to search for 4-SNV interactions. Unlike BOOST and MPI3SNP, which code each bi-allelic SNV to 3 bit-vectors, our algorithm uses 1 bit-vector to store each SNV, enabling more efficient use of modern CPUs. Note that similar to all other exhaustive search algorithms, BitEpi tests all possible combinations of SNVs. BitEpi does not reduce the number of *m*-SNV combinations but the time spent at each test. Furthermore, BitEpi uses entropy statistics, which has been demonstrated to better fit sparse contingency tables in epistasis analysis [22, 23, 19]. We also provide a Python program that computes *p*-value for the statistics used in BitEpi.

As polygenic diseases may be driven by large numbers of individual SNVs as well as interactive SNVs, we developed EpiExplorer to visualize this interplay. EpiExplorer translates a list of interactions into a dynamic Cytoscape [24] graph. It offers a graphical user interface for ease of use and can perform various filtering and highlighting on the graph. Visual elements such as node shape, colours and size are used to represent different genomic or statistic features, such as SNV annotations or the interaction effect size.

## 2 Material and Methods

Processing each combination of SNVs includes two steps: the counting step to find the frequency of genotype combinations and the power analysis to compute the association power and the interaction effect size. The counting step is responsible for most of the execution time. Section 2.1 describes a bitwise process to speed up computing the contingency table for up to four SNVs. The accuracy to identify true epistatic interactions depends on the method used for power analysis. The statistics used to evaluate association power and the interaction effect size from the contingency table are then described in Section 2.2. Section 2.3 describes how *p*-values are computed for the statistics used in BitEpi and Section 2.4 explains the features of EpiExplorer. We elaborate on our experimental setup in Section 2.5.

### 2.1 Counting

The input to BitEpi is a set of bi-allelic SNVs where there are three possible genotypes (0/0, 0/1 and 1/1). Multi-allelic SNVs should be broken into multiple biallelic SNVs before the analysis (i.e. using bcftools norm [25]). Given *m* is the order of the analysis (number of interactive SNVs), the size of the contingency table is 3*^m^* rows and two columns. Each row represents a different genotype combination for the selected SNVs. Columns represent the case and the control cohorts. Each entry of the table is the number of samples with a specific genotype for the selected SNVs in the case or control cohort. Table 1 illustrates an example contingency table for a pair of SNVs: A and B. The fifth row of the table explains that there are 34 cases and 46 controls with a heterozygous genotype for both A and B.

**Table 1:** An example contingency table for 2-SNV interaction of two SNVs: A and B.

| A | B | Case | Control |
| --- | --- | --- | --- |
| 0/0 | 0/0 | 23 | 424 |
| 0/0 | 0/1 | 263 | 423 |
| 0/0 | 1/1 | 534 | 634 |
| 0/1 | 0/0 | 87 | 45 |
| 0/1 | 0/1 | 34 | 46 |
| 0/1 | 1/1 | 56 | 345 |
| 1/1 | 0/0 | 345 | 34 |
| 1/1 | 0/1 | 56 | 64 |
| 1/1 | 1/1 | 456 | 547 |

To speed up the process of counting samples in each cohort with the same genotype, we have implemented a fast bitwise algorithm. Bitwise representation of genotypes allows the genotypes of multiple samples to be stored in a machine word (64-bit) and processed in an operation (bit-level parallelization). In our algorithm, a genotype is encoded using two bits (i.e. 00, 01 and 10 for 0/0, 0/1 and 1/1 respectively) and stored in a byte (8-bits). The remaining 6 bits are set to 0. Thus, 8 samples can be stored in a 64-bit machine word (the parallelization factor is 8 samples per operation). Each SNV is stored in 1 bit-vector (**1-Vector** bitwise approach). Our algorithm uses bitwise SHIFT and OR operators to combine genotypes of up to 4 SNVs. In the resulting vector, each byte represents the genotype of all *m* SNVs for a sample. Thus, the counting process loops through the resulting vector and counts the frequency of each byte.

Since 2-bit encoded genotypes are combined in an 8-bit, the algorithm is limited to combining a maximum of 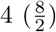 SNVs. It is possible to modify the implementation such that it combines 8 genotypes in a 16-bit machine word (2 bytes) but this would double the theoretical algorithm complexity because the resulting vector would be double the length and a linear reading of it would take twice as long. The current solution hence represents the optimal trade-off between speed and complexity.

Figure 1 is an example that shows the binary representation of genotypes of four different SNVs: A, B, C and D across 8 samples (4 cases and 4 controls). The second, third and fourth SNVs are then shifted to the left by 2, 4 and 6 bits respectively. Next, all four SNVs are combined using bit-wise OR operations. These two steps are also shown in Figure 1. In the resulting array, each byte represents a genotype combination for a sample (a row in the contingency table). For example, 00010010 (for sample S4) represents the row in which D and B have the 0/0 genotype, C has the 0/1 genotype and A has the 1/1 genotype. To form the contingency table, BitEpi loops through the OR vector and counts the occurrences of each byte.

**Figure 1:**
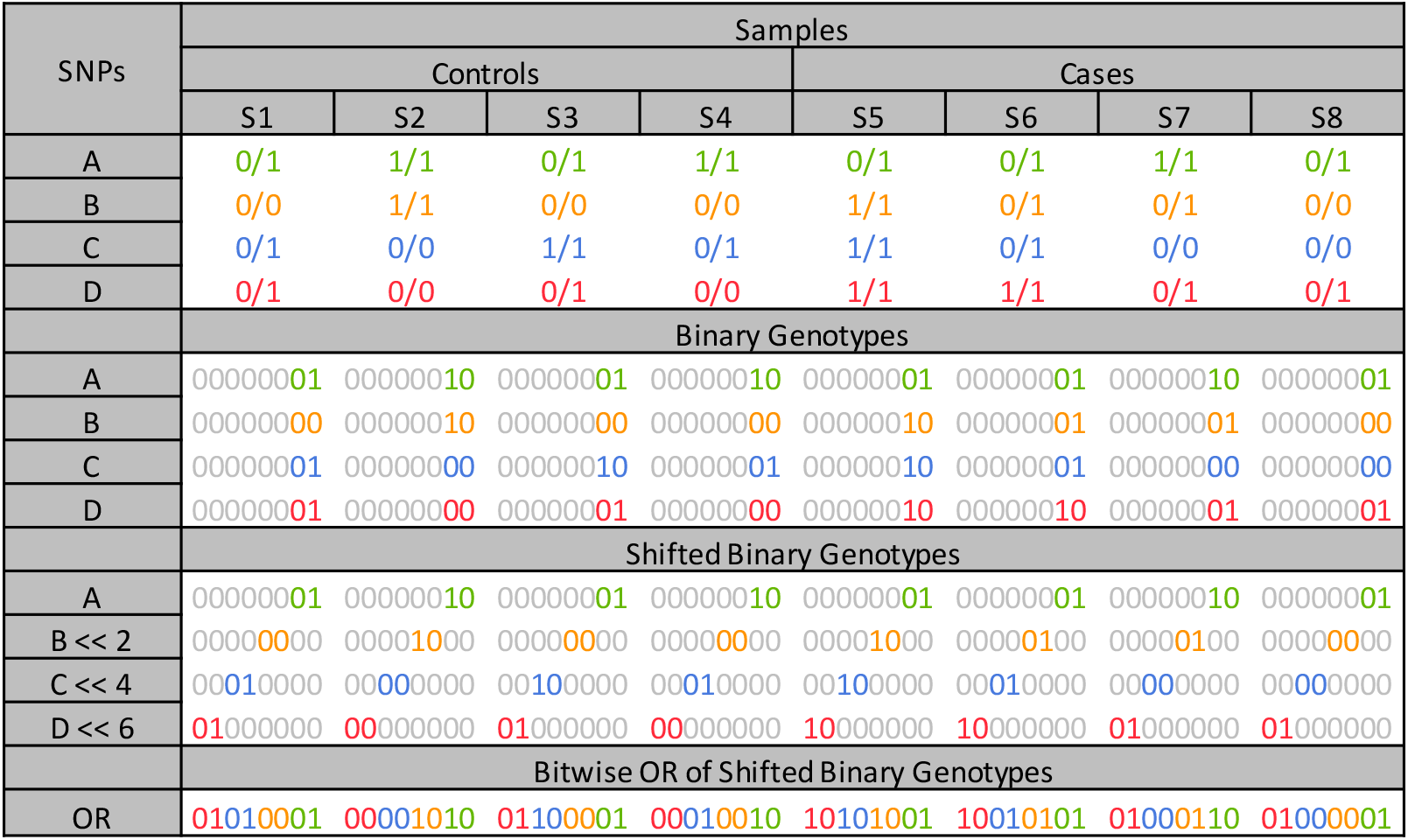
The bitwise representation of 4 example SNVs (A, B, C, and D) and the shifted bit-vectors as well as combined bit-vector.

BitEpi eliminates the shift operations at each test by pre-computing 2, 4 and 6 bit shifted versions of the entire dataset (producing 3 extra copies) and storing them in memory before the analysis. Since the number of SNVs for an exhaustive epistasis analysis is limited, the redundancy in memory usage and the time to precompute shifted datasets are negligible.

Our 1-Vector bitwise approach is different from the 3-Vector bitwise approach used in BOOST and MPI3SNP. Algorithm 1 and Algorithm 2 illustrates the 3-Vector and 1-Vector bitwise approaches to compute one column of the contingency table in a *m*-SNVs interaction (i.e case column or control column). Both cohorts can be processed using the same algorithm.

Here, *C* represent a column of the contingency table where *C*[*i*] is the number of samples in the *i*^th^ row of the table (*i* starts from 0).{*P*[1] *… P*[*m*]} represent *m* SNVs and *R* is a temporary variable (a 64-bit machine word).

In Algorithm 1, each SNV is encoded into 3 bit-vectors, *υ*[1], *υ*[2] and *υ*[3]. Each bit-vector corresponds to a genotype (0/0, 0/1 and 1/1 respectively). For *P*[*i*], if the *j*^th^ sample has the 0/1 genotype, then the *j*^th^ bit in *P* [*i*]*.υ*[2] is set to 1. Each bit-vector is stored in an array of 64-bit machine words where each word contains the information for 64 samples (1 bit per sample). Thus the parallelization factor is 64 samples per operation. *P* [*i*]*.υ*[*j*][*k*] represents the *k*^th^ word of the *j*^th^ vector of the *i*^th^ SNV. There are 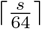 words in each vector where *s* is the number of samples in the cohort (i.e. cases or controls). The core operation of Algorithm 1 includes *m* bit-wise AND operations, a *BitCount* operation to count number of set bits (1’s) in the result of AND operations (*R*) as well as an ADD operation. In this program there are *m* nested loops each iterating from 1 to 3. *x_i_* is the iterator for the *i*^th^ loop. These loops result in the complexity of 3*^m^s* to perform each test.

In contrast, our proposed 1-Vector bitwise method shown in Algorithm 2 does not have the 3*^m^* exponential complexity The downside of this is a lower parallelization factor (8 compared to 64). In Algorithm 2, *P^k^*[*i*]*.υ*[*j*] represents the *j*^th^ word in the bit-vector of *i*^th^ SNV shifted *k* bits to the left. The core operation of the algorithm is *m* bitwise OR operation and 8 increment operations. *R.byte*[*b*] represents *b*^th^ byte in *R* (*R* consist of 8 bytes).

While lower bit-level parallelisation slows down BitEpi’s counting algorithm (Algorithm 2) compared to BOOST’s and MPI3SNP’s counting algorithm (Algorithm 1), the absence of the exponential component (3*^m^*) in complexity of Algorithm 2 makes it overall faster than Algorithm 1, for *m* > 2 (as evident by the experimental results Table 2).

**Algorithm 1:**
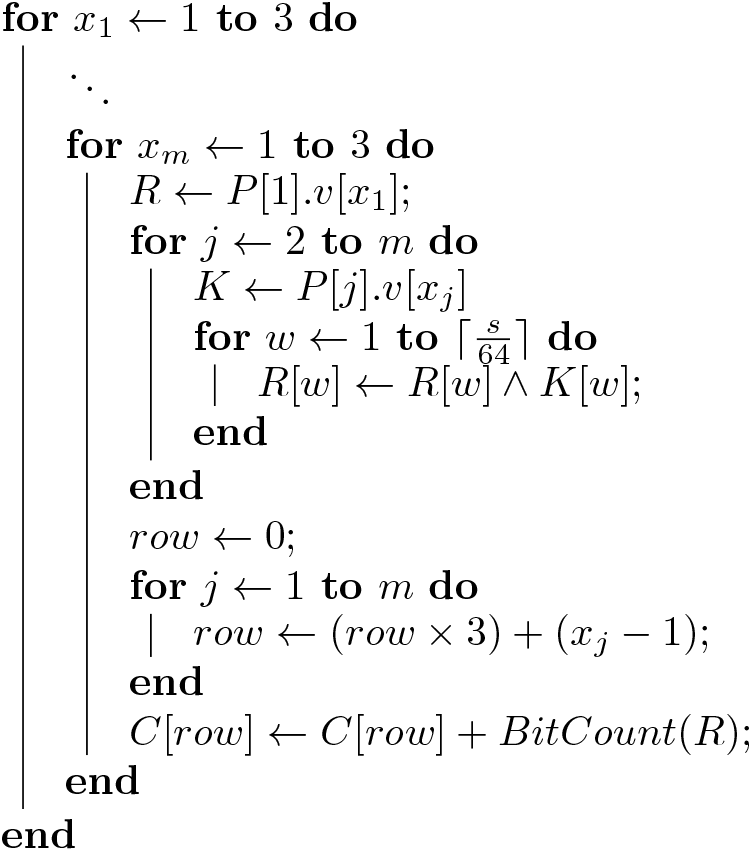
3-Vector bitwise algorithm used in BOOST and MPI3SNP

**Algorithm 2:**
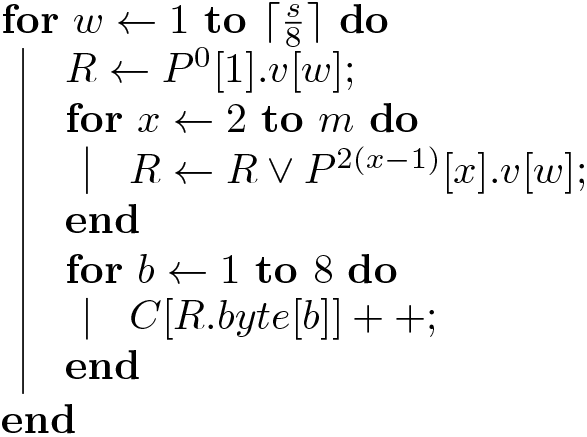
1-Vector bitwise algorithm used in BitEpi

**Table 2:** The execution time (in seconds) of epistasis algorithms for 2,000 samples and different numbers of SNVs. The process is killed if it takes more than an hour to complete and the execution time is not measured. If the execution time is less than a second it is reported as 1 in this table. All programs are executed with 16 parallel threads. Highlighted execution times are used to compute the average test time (see Figure 3).

| Order of Epistasis | Algorithm | Number of SNVs |  |  |  |  |  |  |  |  |
| --- | --- | --- | --- | --- | --- | --- | --- | --- | --- | --- |
|  |  | 100 | 200 | 500 | 1,000 | 2,000 | 5,000 | 10,000 | 20,000 | 50,000 |
| 2-SNV | BitEpi | 1 | 1 | 1 | 1 | 1 | 4 | 11 | 40 | **227** |
|  | BOOST | 1 | 1 | 1 | 1 | 1 | 1 | 4 | 15 | **56** |
|  | MDR | 1 | 1 | 2 | 6 | 29 | 148 | 390 | **2619** | - |
| 3-SNV | BitEpi | 1 | 1 | 4 | 28 | **221** | - | - | - | - |
|  | MPI3SNP | 1 | 1 | 6 | 47 | **375** | - | - | - | - |
|  | MDR | 3 | 14 | 212 | **2121** | - | - | - | - | - |
| 4-SNV | BitEpi | 1 | 15 | **460** | - | - | - | - | - | - |
|  | MDR | 51 | **834** | - | - | - | - | - | - | - |

### 2.2 Statistics

BitEpi computes two metrics for each combination of SNVs: the combined association power (*β*) and the interaction effect size (*α*). While *α* precisely identifies the interaction between SNVs, *β* is needed to compute *α*.

*β* is an entropy metric designed based on the concept of set-purity in the Gini-Index. The purity of a set *p* is computed using Equation 1 where *x* and *y* represent the number of case and control samples in the set. Each row of the contingency table represents a set of samples. The weighted average purity of these sets represents the combined association power of the given contingency table (*β*). The weight for each set is the ratio of the number of samples in the set to the total number of samples. Assuming *x_i_* and *y_i_* represent the number of case and control samples in the *i*^th^ row of the contingency table, the combined association power is computed using Equation 2 where 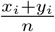 and 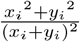 are weight and purity of *i*^th^ set (row) respectively.

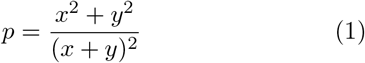

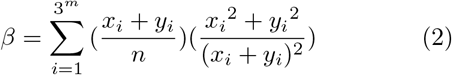

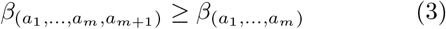

A high combined association power of a set of SNVs does not necessarily indicate a strong interaction between those SNVs. Note that the combined association power is always greater or equal when adding an SNV to the set (Equation 3 proven in the end of *Supplementary Data*). For example, in the left-most graph of Figure 2, *β_AB_*, *β_BC_* and *β_ABC_* are all high. However, they are all driven by the high value of *β_B_*. There is no strong interaction between A and B or B and C as combining B with A or C only slightly increases the association power of SNV B. In a similar fashion combination of ABC slightly increases the association power of BC (no strong 3-SNV interactions).

**Figure 2:**
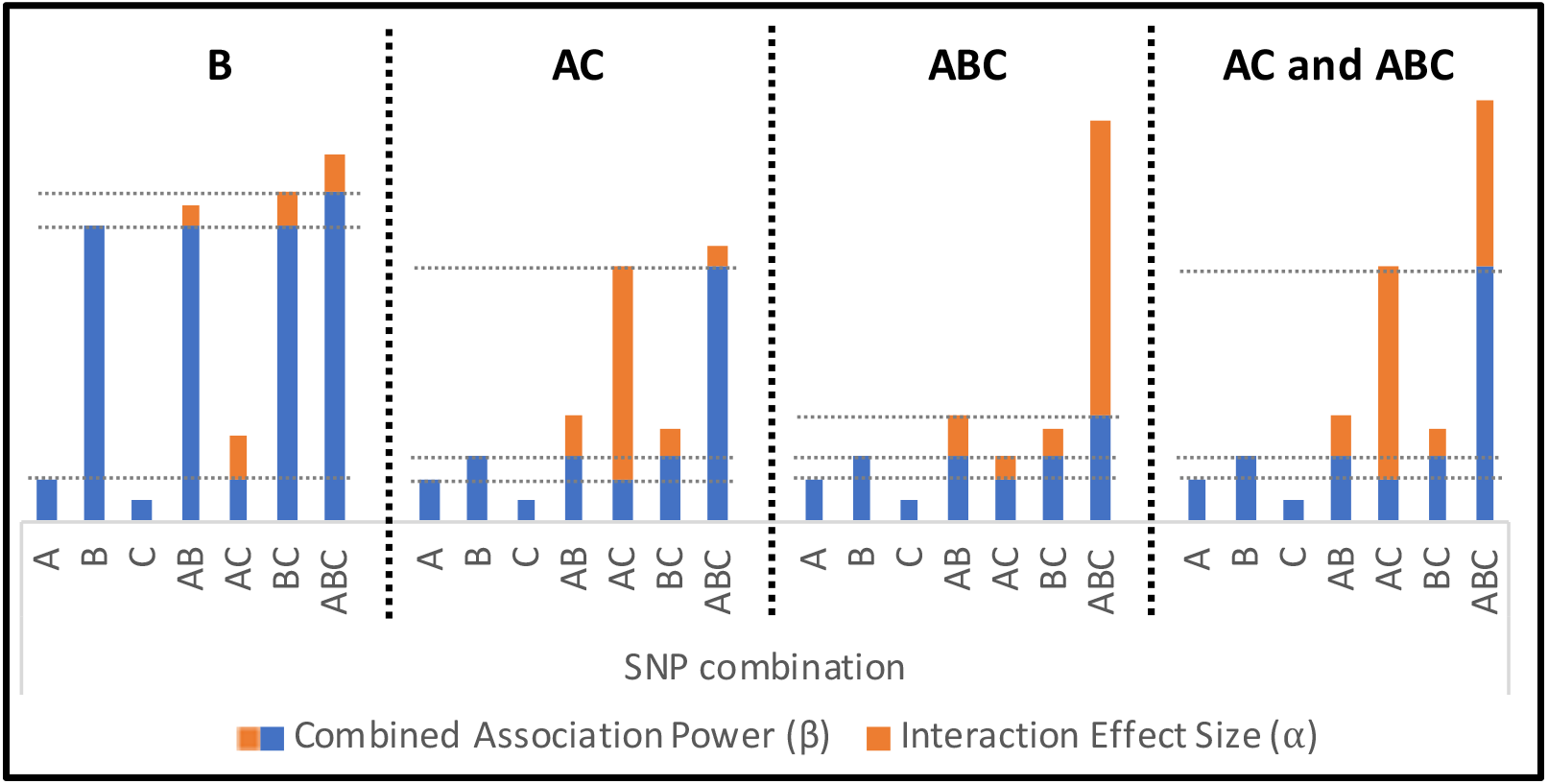
Four examples illustrating the effects of SNVs A, B and C. Bar height (blue and orange together) represent the maximum association power (*β*), while orange visualizes the interactional component (*α*) of the total association. From left to right: **(B)** SNV B strongly associated with the phenotype and none of the 2-SNV and 3-SNV combinations add considerably to the association power of SNV B. In this case, there is no interaction. **(AC)** Neither of SNVs shows strong association power but the combination of AC increases the association power of SNV A significantly. Adding B to the pair of AC has a minor effect on the association power. In this case, A and C strongly interact with each other. **(ABC)** Neither of SNVs shows strong association power. Also, 2-SNV combinations do not increase the association power of SNVs considerably. However, a combination of ABC leads to a significant increase in the association power. In this case, there is a strong 3-SNV interaction of ABC. **(AC and ABC)** Neither of SNVs shows strong association power but the combination of AC increases the association power of SNV A significantly. Adding B to the pair of AC further increases the association power of AC. In this case, there is a strong 2-SNV interaction of AC and a strong 3-SNV interaction of ABC.

In epistasis analysis, we are interested in the set of SNVs where the association power is driven by the interaction between all SNVs in the set, as opposed to an individual SNV or through additive effects of SNV subsets. Thus in computing *α*, we look at the gain in the association power that presents only when considering all SNVs together. For example, to compute *α_ABC_*, we subtract from *β_ABC_*, the maximum combined association power of any subset of (A,B,C). Since we know that max(*β_AB_, β_AC_, β_BC_* is greater or equal to max(*β_A_, β_B_, β_C_*, we only need to compute the former. In general, to compute *α* for a *m*-SNV interaction, we should find the maximum *β* of all *m* – 1-SNV combinations.

To formulate *α* computation, assume *G^m^* is a set of *m* SNVs (*a*_1_*, a*_2_, …, *a_m_*) and 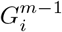 is *G^m^* excluding *a_i_* (i.e. 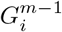 is a subset of *G^m^*). Then, 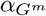 is computed using Equation 4. BitEpi computes *α* and *β* for individual SNVs too (normal GWAS). To compute *α* for an individual SNV 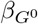 is computed as the purity of the set that includes all samples.

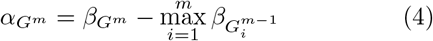

In order to compute 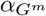, the program needs to compute 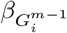. Since there could be common SNVs between two sets of *m* SNVs, the same 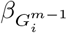 should be recomputed multiple times. For example, to compute *α*_(*A,B,C,x*)_ where *x* could be any SNV in the dataset other than A, B and C, *β*_(*A,B,C*)_ should be recomputed. This results in a huge computational redundancy. To avoid this redundancy, prior to computing 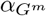, BitEpi computes all lower-order 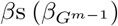 and stores them in a multi-dimensional array. Using a multi-dimensional array to store *β* for all possible (*m* – 1)-SNV combinations results in memory redundancy (memory is allocated but not used). However, lower order *β* values are accessed frequently and a multi-dimensional array allows for the fastest retrieval.

BitEpi can perform any combination of *m*-SNV *α* and *β* test in the same analysis where *m* could be 1, 2, 3 or 4. There is a special mode of operation called best. For each SNV, the best mode lists the 2-SNV, 3-SNV and 4-SNV interaction with the highest *α*. The user can choose to list the significant interactions with the highest *α* or *β*. This can be done by identifying either the number of top interactions to be reported or a threshold on *α* or *β* (all interactions that exceed the threshold will be reported).

BitEpi is implemented in `C++` with multi-threading. Each SNV combination is independent of other SNV combinations, thus SNV combinations can be processed on different machines or processors. We currently implement an efficient multi-threading that balances the number of SNV combinations to be processed on each processor of a High-performance compute node.

It includes a Python wrapper so that it can be installed using pip and used in a Python program. An R script is provided to turn BitEpi best output to a static igraph graph and a dynamic Cytoscape graph.

### 2.3 *p*-value calculation

The BitEpi code-base includes a Python program that computes *p*-values for the given set of interactions. Since the underlying distribution for the *α* and *β* statistics are unknown, we create the Null distribution empirically for each SNV or SNV interaction. This is done by permuting the phenotype many times and computing *α* and *β* value on the permuted data to create a discrete distribution. An empirical *p*-value can be calculated by counting how often the value for the permuted data is equal or larger than the value on the real data and divide it by the number of permutations. For example, if we permute the phenotype 1000 times and in 20 instances the computed *α* is equal or greater than the *α* computed for the actual phenotype, then the *p*-value is 20/1000 = 0.02. However, this approach is not suitable for the small *p*-values we expect. Instead of calculating the simple ratio, we fit a continuous probability distribution to the discrete distribution and compute the *p*-value as 1 *CDF* (*observation*), where CDF is the Cumulative Distribution Function. Note that we use the one-way statistics since the only greater value of *α* and *β* are considered extreme.

### 2.4 Visualization

EpiExplorer is able to visualize large numbers of SNV interactions as well as individual SNV associations. It takes lists of interactions and their statistics along with the genomic annotations of the SNVs involved in the interactions. Together, they are transformed into a Cytoscape graph. Cytoscape is a visualization platform designed to work with large complex graphs such as the complete set of epistatic interactions of complex genomic phenotypes. It provides useful functionalities such as various layout (placement of nodes) algorithms as well as styling tools. Visual elements of nodes (shape, colour, size) and edges (colour, thickness) can be used to represent different genomics or statistical features of SNVs or the interaction between them. For example, the size of the node can represent the interaction effect size and the shape of the node can represent if the SNV is in a protein-coding region or not.

EpiExplorer uses Cytoscape Python API to take control of the plot and change the style of it. More importantly, it provides incremental filtering and highlights a feature that accepts complex queries while keeping the layout fixed. For example, all non-coding SNV nodes in the graph can be hidden, while making all micro-RNA SNV nodes visible (put them back to graph). Visualizing them relative to all other nodes, which will remain in the same place, allows for investigating the interaction of micro-RNA and protein-coding genes. From there, SNVs of a specific chromosome can be highlighted by greying out the rest from the graph to produce a publication-ready figure. Note that, such a complex query is only available through EpiExplorer but not Cytoscape. Though this capability is enhanced by Cytoscape offering a range of different layout algorithms, such as SNVs grouped by chromosomes, or the functionality of the region they are located at (i.e. Exone, Intron, Gene name, MIR etc), sorted based on their genomic location or their relevance to the phenotype.

Besides the obvious way of representing interactions where SNVs are nodes and interactions are edges, EpiExplorer offers a second visualization specifically geared towards higher-order interactions. In this second mode, both SNVs and interactions are represented as nodes of the graph. Each interaction node is connected to all the corresponding SNV nodes. This mode is suitable for a more in-depth study focusing on a smaller subset of interactions.

### 2.5 Experimental setup

Several synthetic datasets are used to evaluate the performance and accuracy of BitEpi and compare it with BOOST, MPI3SNP, and MDR.

To test the accuracy (detection power), we use GAMETES [16] to generate ground truth datasets (where the interactive SNVs are known). We create 10 simulated 2-SNV epistasis models with different heritability and Minor Allele Frequency (MAF=0.01 and 0.5) of the interactive SNVs (Pairwise Models: PM1~PM10), see Supplemental Table 4. Each model includes one 2-SNV interaction. We also create 9 epistasis models (Triplet Models: TM1~TM9) each of which includes one 3-SNV interactions (see Supplemental Table 5). For each model, 100 datasets are generated each with 100 SNVs and 2,000 samples (1,000 cases and 1,000 controls). To compute the detection power of an algorithm (A) for a model (M), we process all 100 datasets generated from model M using algorithm A and count how many times the known interactive SNVs are ranked first(i.e. reported to have the highest association power). Any interactions ranked above the known interaction (i.e. reported with an even higher association power) is considered as false positive. Model files with detailed model parameters are available in *Supplementary Data*.

To test execution time, we create much larger datasets by randomly assigning genotypes and phenotypes to samples. Each dataset consists of a different number of SNVs and samples (see Supplemental Table 1, Supplemental Table 2 and Supplemental Table 3 as well as Table 2).

To benchmark the performance of BitEpi against existing tools and test a wider range of epistatic models, we also compare on previously published synthetic datasets [26]. These datasets include 12 Marginal Effect (ME1~ME12) and 40 No Marginal Effect (NME1~NME40) epistasis models where each model includes one 2-SNV interaction. For each epistasis model, 100 datasets each with 100 SNVs and 1,600 samples (800 case and 800 controls) are simulated.

All tests were performed on a machine with dual 10 core Intel Xeon E5-2660 V3 processors running at 2.6 GHz with 25 MB cache and 128 GiB of memory with SUSE Linux Enterprise Server 12 SP4. To compile BitEpi we used gcc version 4.8.5 and glibc version 2.22, no additional libraries were used.

### 2.6 Wellcome Trust Case Control Consortium

To test BitEpi on real datasets, we compare the performance against BOOST on genomic data from the Wellcome Trust Case Control Consortium [27] (WTCCC). For this comparison, we perform an exhaustive search for pairwise interaction in seven case/control datasets (type 1 diabetes, type 2 diabetes, rheumatoid arthritis, inflammatory bowel disease, bipolar disorder, hypertension, coronary artery disease). Each dataset consists of two control cohorts (National Blood Service and British Born in 1958) and one case cohort. In the data preparation, SNPs with Minor Allele Frequencies (MAF less than 5%) and in Linkage-Disequilibrium (LD with *r*^2^ = 0.2) are removed. Outlier samples detected by principal component analysis are also removed. The list of samples (~4900 per dataset) and SNPs (~87, 000 per dataset) used are provided in *Supplementary Data File* (Plink [28] bim and fam file format).

For comparing BOOST and BitEpi on this dataset, we follow the recommended Plink epistasis pipeline for calculating *p*-values, which starts with generating boost data by processing the complete dataset with `--fast-epistasis` (BOOST) and then listing SNPs that are involved in significant interactions (i.e. top 1000 interactions). Then `--epistasis` analysis is performed to compute Logistic-Regression p-value for all the pairs. Note that Logistic-Regression analysis is very slow and cannot be applied to a large number of SNPs, despite the multi-threaded implementation of BOOST (Plink v1.9 epistasis fast [28]). As illustrated in Supplemental Figure 3, to calculate p-values for BitEpi we replace the BOOST score with BitEpi pairwise *α* analysis. We then compare the resulting p-values from the Logistic-Regression analysis at the end of both pipelines.

## 3 Results

### 3.1 BitEpi is faster for higher-order interactions

We compare the execution time of BitEpi’s *α* test against the other state-of-the-art exhaustive epistasis search algorithms, BOOST, MPI3SNP and MDR. Note that BOOST and MPI3SNP perform 2-SNV and 3-SNV analysis respectively, while MDR is the only other method besides BitEpi to process different order of interactions.

As shown in Table 2, BitEpi performs the fastest out of all surveyed methods, this is because the 1-Vector bitwise method does not have exponential complexity (See Methods 2.1). Specifically, BitEpi performs up to 1.7 times faster than MPI3SNP for 3-SNV searches (2,000 SNVs dataset) and up to 65, 76 and 56 times faster than MDR for 2-SNV, 3-SNV and 4-SNV searches (20,000 SNVs, 1,000 SNVs, and 200 SNVs datasets), respectively. It is scalable to the largest dataset (50,000 SNVs), with BOOST the only other method to also achieve this. Here, BOOST’s specialized 2-SNV algorithm is up to 4 times faster than BitEpi on this specific use case. Note that we report the largest dataset the algorithms were capable of processing within the given compute resources and time-cutoff (1 hour).

BitEpi’s observed speedup over MPI3SNP is because the 1-Vector algorithm in BitEpi is independent of the order of the epistasis interaction. This allows BitEpi to perform the individual interaction tests at the same speed, irrespective of whether a 2-SNV, 3-SNV or 4-SNV interaction is tested. To quantify the improvement, we compute the test time for each order. As the order of epistasis increases, the number of tests that need to be performed also increases. We hence normalize execution time by the number of tests performed to be able to directly compare the individual 2-SNV, 3-SNV and 4-SNV tests between 3-Vector and 1-Vector bitwise algorithms. We compute the average test time as 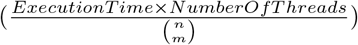 Where 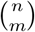 is the number of *m*-SNV tests in a dataset with *n* SNVs for the datasets highlighted in Table 2.

Figure 3 shows that the 1-Vector approach used in BitEpi can keep the execution time constant (2.7~2.9*μs*) for all orders tested. Please note, the reported execution time per test comprises the construction of the contingency table (first step) as well as performing the statistical test on the contingency table (second step). While the 1-Vector algorithm keeps the execution time of the first step constant, the complexity of the statistical test in the second step is exponential with the number of SNVs in the interaction (as it increases the number of rows in contingency table). However, the statistical test is executed on a small contingency table while the first step has to processes a large array of genotypes for thousands of samples. The influence of the exponential component on the overall runtime is hence negligible.

**Figure 3:**
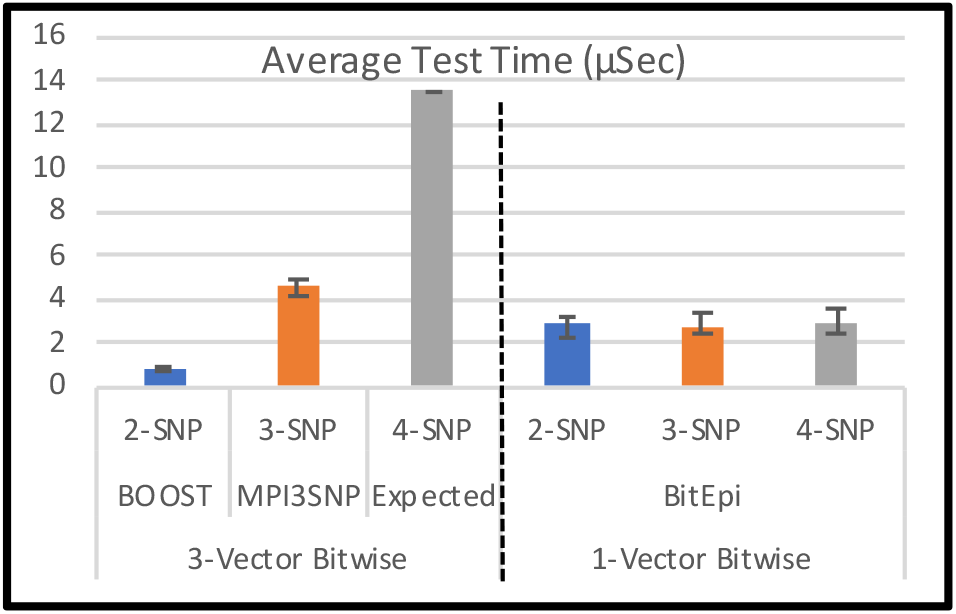
The average runtime per test for 2,3, and 4-SNP interactions, comparing the 3-Vector bitwise approach (left) with BitEpi’s 1-Vector bitwise approach (right). The expected 4-SNV average test time with the 3-Vector bitwise approach is computed as *MPI*3*SNP* × 3. The average test time is computed based on the highlighted execution time in Table 2. Error bars represent standard error.

BitEpi has a 1.7 (3-SNV) and 4.7 (4-SNV) fold speedup compared to the 3-Vector method used in BOOST and MPI3SNP. Specifically, the 2-SNV test in BOOST takes 0.7*μs* on average, while the 3-SNV test in MPI3SNP takes 4.5*μs* on average. As there are no 4-SNV bitwise methods published to date, we extrapolate from the 3-SNV searches resulting in execution time of 4.5*μs* × 3 = 13.5*μs*. The complexity of the 3-Vector bitwise method grows exponentially with the number of interactive SNVs (3*^m^*). For completeness, we list the experimentally determined runtime for the only other 4-SNV method, MRD, which does not use bitwise procedures, which is substantially slower in all categories.

BitEpi also scales linearly with the number of samples, as shown in Supplemental Table 2 (total execution time with increase samples) and Supplemental Figure 1.b (normalized by samples). However, BitEpi scales exponentially with the number of SNVs, *υ*, resulting in *O*(*υ*^2^), *O*(*υ*^3^) and *O*(*υ*^4^) for 2-SNV, 3-SNV and 4-SNV, respectively, shown in Supplemental Table 1 (total execution time) and Supplemental Figure 1.a (normalized execution time per SNV). Both execution times can be curbed by parallelization, as shown in Supplemental Table 3 and Supplemental Figure 1.c, in which using 2,4,8 and 16 CPUs results in a non-saturated, near-linear speed-up.

### 3.2 BitEpi is more accurate in detecting interactions

To compare the accuracy of BitEpi with BOOST and MPI3SNP, we compute the detection power for all models simulated by GAMETES [16] including models simulated by others [26]. Figure 4a shows the 2-SNV detection power of BitEpi and BOOST for the models we simulated. Except for the Pairwise-model 1 (PM1) where both methods result in poor detection power, BitEpi performs better than BOOST (i.e. between 1.22 and 1.33 times more accurate) and reaches 100% detection power for PM7~PM10 Models.

**Figure 4:**
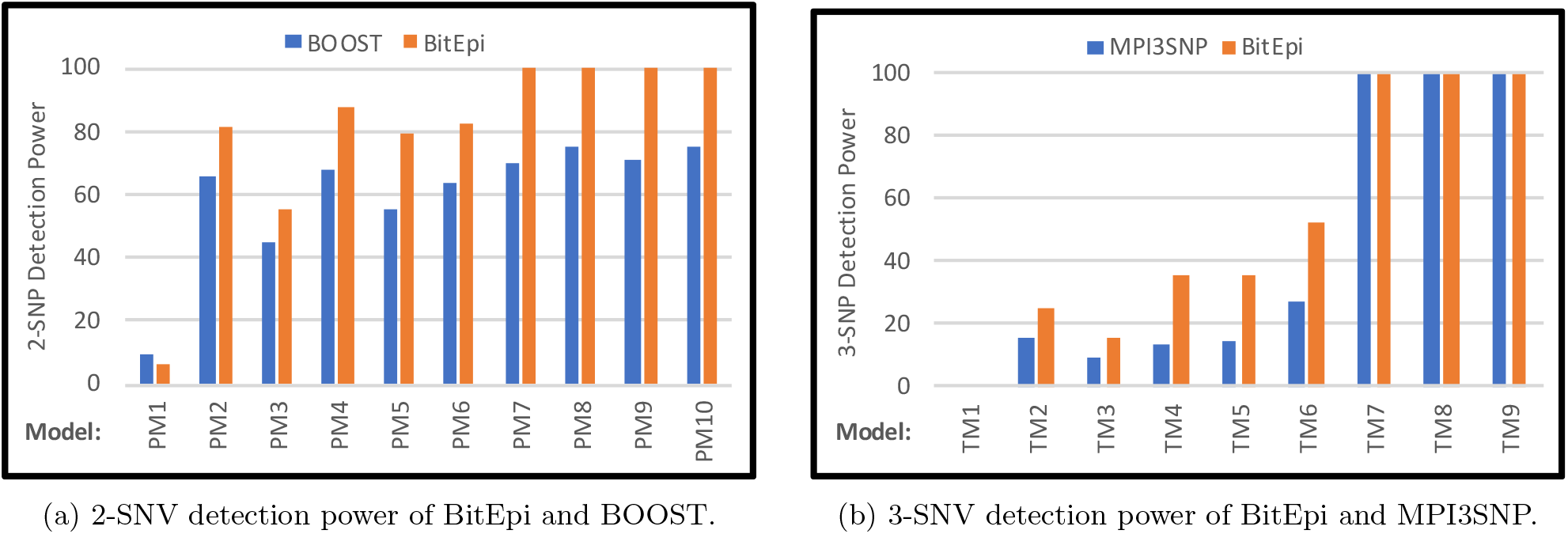
Compare detection power of BitEpi with BOOST and MPI3SNP for 2-SNV and 3-SNV analysis.

Figure 4b shows the 3-SNV detection power of BitEpi and MPI3SNP. BitEpi performs better than MPI3SNP for Triplet-models TM2~TM6 Models (i.e. between 1.56 and 2.09 times more accurate), and equivalent for the rest. Numerical comparisons are available in Supplemental Table 4 (2-SNV) and Supplemental Table 5 (3-SNV).

We also compute the 2-SNV detection power of BitEpi and BOOST for 12 ME (Marginal Effect) and 40 NME (No Marginal Effect) epistasis models simulated in [26]. Supplemental Figure 2 illustrates the comparison result. Numerical comparisons are available in Supplemental Table 6 and Supplemental Table 7. BitEpi’s detection power for ME models is 44% higher than BOOST’s on average. For NME models, BitEpi’s average detection power is the same as BOOST’s.

Out of the 71 epistasis models we have evaluated, BitEpi performs better than other methods in 24 cases, similar to other methods in 39 cases and is less accurate than other methods in 8 cases. This indicates that due to the accurate isolation of interaction effect sizes, BitEpi eliminates false positives more effectively.

### 3.3 p-value and Visualization

To demonstrate the capability of our *p*-value calculation and visualization program (EpiExplorer) we created a synthetic dataset using GAMETES where the phenotype is a function of three truth variables: an individual SNV (A), a 2-SNV (B-C) and a 3-SNV (D-E-F) interactions (E123 Dataset). We have processed the dataset with BitEpi to list all significant 1,2, and 3-SNV associations. Figure 5a shows the *α* statistic for the top 10 variables of each order (sorted by *α*) as well as the *α* statistic of 10 randomly selected variables for each order as a comparison. The highlighted bar represent the truth variable for each order.

**Figure 5:**
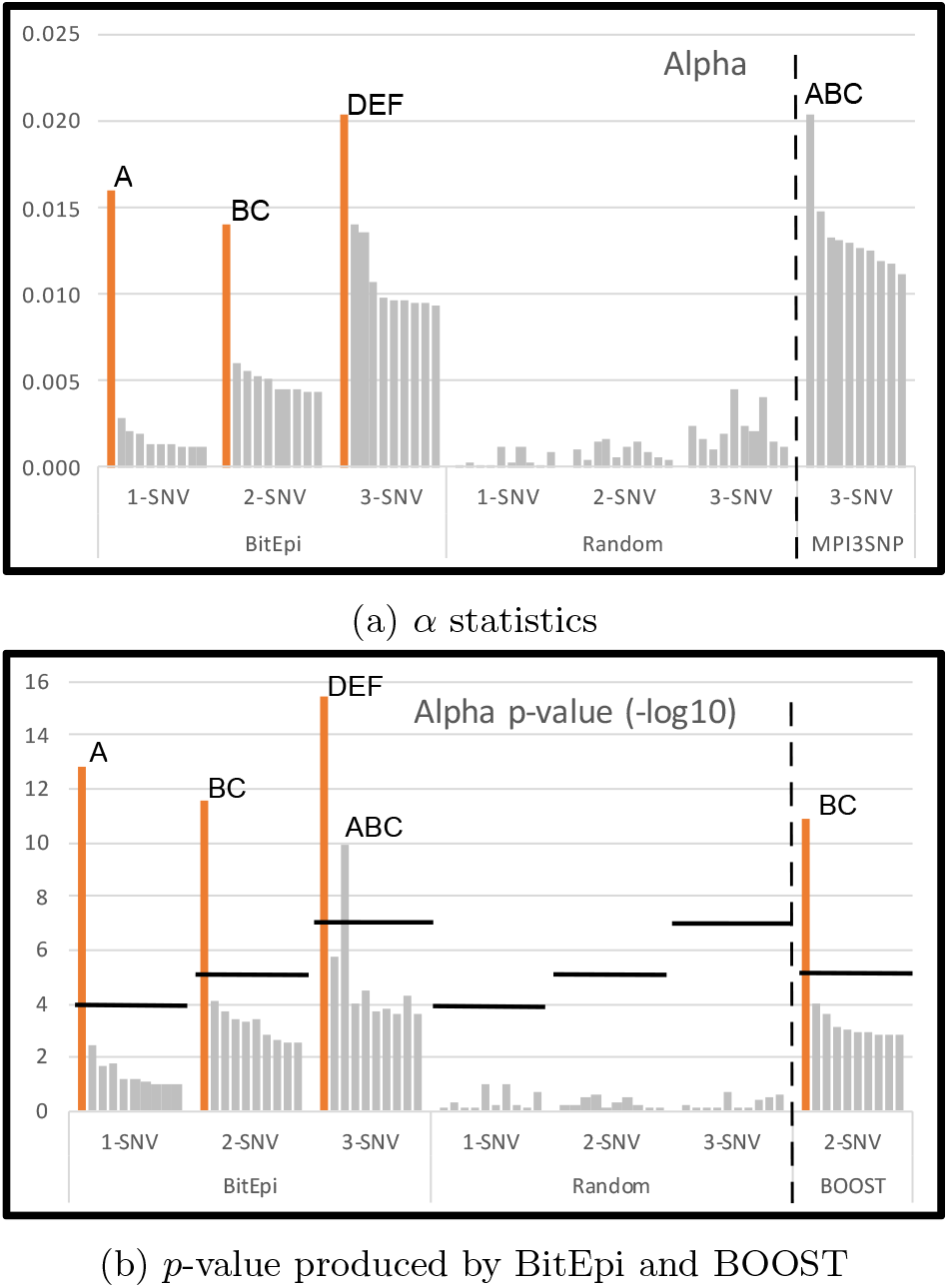
*α* and *p*-value for the top 10 variables and 10 randomly selected variables of E123 dataset.

We next calculated the *p*-values, to quantify the significance of the clear separation seen between the *α* statistics of the truth variables and the noise. We plot the –*log*10 transformed *p*-value in Figure 5b. The significance threshold 0.05 was set and *p*-values corrected for multiple testing using Bonferroni correction with 100, 4,950, and 161,700 tests, respectively. This is because there are 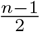 times more 2-SNV combinations than 1-SNVs combinations, where *n* is the number of SNVs, and 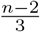 times more 3-SNV combinations compared to 2-SNV combinations.

Among the top 10 evaluated interactions, only the truth variable is statistically significant for 1-SNV and 2-SNV. For the 3-SNV interaction, another variable crossed the significance threshold (ABC), besides the true interaction between DEF. Upon closer inspection, we determined this to be an artifact in the simulated data, where only the truth variables are modelled explicitly. As a result, other interactions can have an association with the phenotype by random chance and the probability of this occurring increases with the number of SNV combinations that are included in the dataset, i.e. 161,700 3-SNV combination when there are only 1000 SNV in the dataset.

To benchmark BitEpi against other state-of-the-art tools, we also process the E123 dataset with BOOST and MPI3SNP. BOOST’s *p*-values for the top 10 2-SNV interactions are similar to those produced by BitEpi (Figure 5b). However, MPI3SNP’s entropy-based method incorrectly detects the combination of A, B and C as the strongest triplet interaction (Figure 5a). This is the same combination that also crossed the threshold in BitEpi’s *p*-value calculation. As explained, it may have a weak association by accident, but should not have been prioritized over the explicitly modeled DEF interaction.

For comparison, we also show the *α* and *p*-values for 10 randomly selected variables of each category, demonstrating the noise level of the data.

We use EpiExplorer to provide a visual representation of Figure 5a, highlighting the identified interactions. As discussed, the corresponding dataset contains all 1-SNV, 2-SNV and 3-SNV interactions with A, BC, and DEF as the respective truth variables in each category. In Figure 6, we plot the top 5 interactions in each category as determined by *α*, as well as the other SNV that are part of the interactions, resulting in 5 blue triangles (3-SNV), 5 red diamonds (2-SNV) and 5 larger green circles (1-SNV), as well as 14 other involved 1-SNV interactions (small green circles). The interaction with the biggest *α* according to Figure 5a is between DEF and is also visualized as the largest item in Figure 6. The second and third largest element is A and BC and visualized with a size according to their *α* value. The remaining unlabeled interactions visualize the remaining non-truth interactions in the top 5, which result in markedly smaller graph elements. The exception is the earlier discussed ABC interaction and another 3-SNV interaction, which have a similar *α* value as A and BC (Figure 5a). It is hence important to combine the visualization with the *p*-value calculation to identify interactions of importance in a discovery scenario.

**Figure 6:**
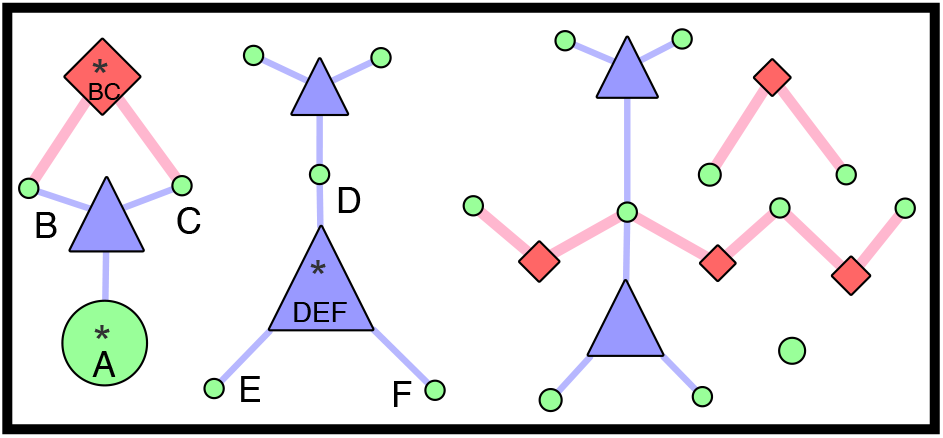
The example plot of the top 5 variables in 1-SNV, 2-SNV and 3-SNV category.

### 3.4 Real Dataset

Demonstrating BitEpi’s capabilities on a real dataset, we compare BitEpi to BOOST on seven case/control datasets from WTCCC. To make the two methods directly comparable, we use BOOST and BitEpi results inside the 2-step Plink-based epistasis framework. We compare the resulting Logistics-Regression *p*-value of the top 100 pairs in each dataset.

Figure 7 shows that both pipelines detect interactions with *p*-value less than 10^−9^. For pairs that are exclusively detected by either pipeline, BitEpi performs slightly better and detects pairs with lower Logistics-Regression *p*-value, which confirms BitEpi’s applicability to real-world data. It also indicates that the *α* score is a reliable proxy for screening large scale datasets.

**Figure 7:**
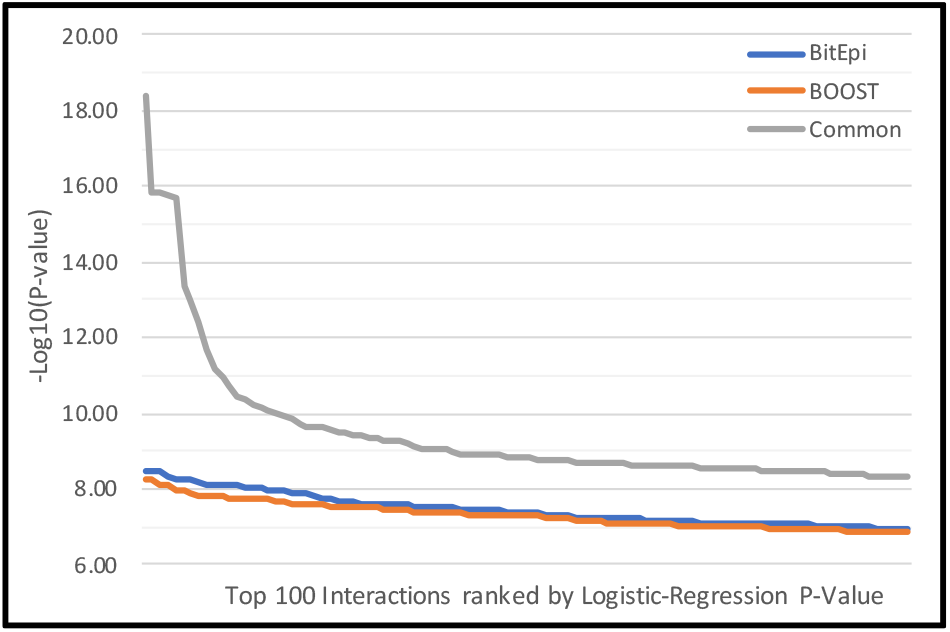
The figure shows the *p*-value of the identified interaction pairs, color-coded by being uniquely discovered by BitEpi, BOOST, or discovered by both.

## Discussion

We demonstrated that the current best practice for exhaustive epistasis search tools (BOOST, MPI3SNP) can be improved upon in both speed and accuracy. While heuristics such as Random Forest remain necessary to reduce the initial search space, BitEpi is then capable of detecting higher-order interactions of up to 4-SNV exhaustively, resulting in an up to 1.7 and 56 fold faster execution time than other surveyed methods for 3-SNV and 4-SNV searches, respectively.

BitEpi uses a novel 1-Vector bitwise approach that is designed for higher-order analysis and allows modern 64-bit machines to be used more effectively than the previous 3-Vector bitwise approaches. It also isolates the interaction effect size using an entropy-based metric to eliminate false positives. BitEpi visualizes the results in an interactive graph that can be dynamically scaled and rearranged, streamlining interpretation and publication.

Future improvements will cover the use of epistatic genomic relationship matrix (EGRM) to control for the effect of diversity [29], as well as more advanced visualization approaches using either d3 or Cytoscape JavaScript library for dynamic web-based visualization. We also plan to add an end-to-end integration with cloud-based Random Forest implementation VariantSpark [15], to enable epistasis search within the ultra-high dimensional data of whole-genome sequencing cohorts.

## Supplementary Data

Supplemental Data includes extended methods and additional figures and tables in SupplementaryData.pdf and addtional information about Wellcome Trust Case Control sample SupplementaryDataFile.tar.gz.

## Acknowledgments

AB, BH and YJ implemented the algorithm and performed experiments. AB and MK implemented the EpiExplorer. AB, NT, and DB conceived the work and wrote most of the paper. CH refined the methodology section. MK refined and added content to the visualization section. All authors have read and approved the manuscript.

## Availability

Codes and data are publicly available on GitHub https://github.com/aehrc/BitEpi and https://github.com/aehrc/EpiExplorer. BitEpi is also avail-able on CodeOcean https://doi.org/10.24433/CO.3671084.v1.

## References

[1] Wen-Hua Wei, Gibran Hemani, and Chris S Haley. Detecting epistasis in human complex traits. Nature Reviews Genetics, 15(11):722, 2014.

[2] Daniel M Weinreich, Yinghong Lan, C Scott Wylie, and Robert B Heckendorn. Should evolutionary geneticists worry about higher-order epistasis? Current opinion in genetics & development, 23(6):700–707, 2013.

[3] Matthew B Taylor and Ian M Ehrenreich. Higher-order genetic interactions and their contribution to complex traits. Trends in genetics, 31(1):34–40, 2015.

[4] Clément Niel, Christine Sinoquet, Christian Dina, and Ghislain Rocheleau. A survey about methods dedicated to epistasis detection. Frontiers in genetics, 6:285, 2015.

[5] Junliang Shang, Junying Zhang, Yan Sun, Dan Liu, Daojun Ye, and Yaling Yin. Performance analysis of novel methods for detecting epistasis. BMC bioinformatics, 12:475, dec 2011.

[6] Li Chen, Guoqiang Yu, David J. Miller, Lei Song, Carl Langefeld, David Herrington, Yongmei Liu, and Yue Wang. A ground truth based comparative study on detecting epistatic SNPs. In 2009 IEEE International Conference on Bioinformatics and Biomedicine Workshop, pages 26–31. IEEE, nov 2009.

[7] Mathieu Emily. A survey of statistical methods for gene-gene interaction in case-control genome-wide association studies. Journal de la Societe Française de Statistique, 159(1):27–67, 2018.

[8] Heather J Cordell. Epistasis: what it means, what it doesn’t mean, and statistical methods to detect it in humans. Human molecular genetics, 11(20):2463–2468, 2002.

[9] Margaret J Eppstein and Paul Haake. Very large scale relieff for genome-wide association analysis. In 2008 IEEE Symposium on Computational Intelligence in Bioinformatics and Computational Biology, pages 112–119. IEEE, 2008.

[10] Makiko Yoshida and Asako Koike. Snpinterforest: a new method for detecting epistatic interactions. BMC bioinformatics, 12(1):469, 2011.

[11] Xia Cao, Guoxian Yu, Jie Liu, Lianyin Jia, and Jun Wang. Clustermi: Detecting high-order snp interactions based on clustering and mutual information. International journal of molecular sciences, 19(8):2267, 2018.

[12] Yan Meng, Qiong Yang, Karen T Cuenco, L Adrienne Cupples, Anita L DeStefano, and Kathryn L Lunetta. Two-stage approach for identifying single-nucleotide polymorphisms associated with rheumatoid arthritis using random forests and bayesian networks. In BMC proceedings, volume 1, page S56. BioMed Central, 2007.

[13] Leo Breiman. Random forests. Machine Learning, 45(1):5–32, Oct 2001.

[14] Rui Jiang, Wanwan Tang, Xuebing Wu, and Wen-hui Fu. A random forest approach to the detection of epistatic interactions in case-control studies. BMC bioinformatics, 10(1):S65, 2009.

[15] Arash Bayat, Piotr Szul, Aidan R O’Brien, Robert Dunne, Oscar J Luo, Yatish Jain, Brendan Hosking, and Denis C Bauer. Variantspark, a random forest machine learning implementation for ultra high dimensional data. bioRxiv, page 702902, 2019.

[16] Ryan J Urbanowicz, Jeff Kiralis, Nicholas A Sinnott-Armstrong, Tamra Heberling, Jonathan M Fisher, and Jason H Moore. Gametes: a fast, direct algorithm for generating pure, strict, epistatic models with random architectures. BioData mining, 5(1):16, 2012.

[17] Xiang Zhang, Shunping Huang, Fei Zou, and Wei Wang. Team: efficient two-locus epistasis tests in human genome-wide association study. Bioinformatics, 26(12):i217–i227, 2010.

[18] Xiang Wan, Can Yang, Qiang Yang, Hong Xue, Xiaodan Fan, Nelson LS Tang, and Weichuan Yu. Boost: A fast approach to detecting gene-gene interactions in genome-wide case-control studies. The American Journal of Human Genetics, 87(3):325–340, 2010.

[19] Junliang Shang, Yingxia Sun, Jin-Xing Liu, Jun-feng Xia, Junying Zhang, and Chun-Hou Zheng. Cinoedv: a co-information based method for detecting and visualizing n-order epistatic interactions. BMC bioinformatics, 17(1):214, 2016.

[20] Jason H Moore and Peter C Andrews. Epistasis analysis using multifactor dimensionality reduction. In Epistasis, pages 301–314. Springer, 2015.

[21] Christian Ponte-Fernández, Jorge González-Domínguez, and María J Martín. Fast search of third-order epistatic interactions on cpu and gpu clusters. The International Journal of High Performance Computing Applications, page 1094342019852128, 2019.

[22] Ting Hu, Yuanzhu Chen, Jeff W Kiralis, Ryan L Collins, Christian Wejse, Giorgio Sirugo, Scott M Williams, and Jason H Moore. An information-gain approach to detecting three-way epistatic interactions in genetic association studies. Journal of the American Medical Informatics Association, 20(4):630–636, 2013.

[23] Sangseob Leem, Hyun-hwan Jeong, Jungseob Lee, Kyubum Wee, and Kyung-Ah Sohn. Fast detection of high-order epistatic interactions in genome-wide association studies using information theoretic measure. Computational biology and chemistry, 50:19–28, 2014.

[24] Paul Shannon, Andrew Markiel, Owen Ozier, Nitin S Baliga, Jonathan T Wang, Daniel Ramage, Nada Amin, Benno Schwikowski, and Trey Ideker. Cytoscape: a software environment for integrated models of biomolecular interaction networks. Genome research, 13(11):2498–2504, 2003.

[25] Heng Li, Bob Handsaker, Alec Wysoker, Tim Fennell, Jue Ruan, Nils Homer, Gabor Marth, Goncalo Abecasis, Richard Durbin, and 1000 Genome Project Data Processing Subgroup. The Sequence Alignment/Map format and SAMtools. Bioinformatics, 25(16):2078–2079, 06 2009.

[26] Peng-Jie Jing and Hong-Bin Shen. Macoed: a multi-objective ant colony optimization algorithm for snp epistasis detection in genome-wide association studies. Bioinformatics, 31(5):634–641, 2014.

[27] Wellcome Trust Case Control Consortium et al. Genome-wide association study of 14,000 cases of seven common diseases and 3,000 shared controls. Nature, 447(7145):661, 2007.

[28] Shaun Purcell, Benjamin Neale, Kathe Todd-Brown, Lori Thomas, Manuel A.R. Ferreira, David Bender, Julian Maller, Pamela Sklar, Paul I.W. de Bakker, Mark J. Daly, and Pak C. Sham. Plink: A tool set for whole-genome association and population-based linkage analyses. The American Journal of Human Genetics, 81(3):559–575, 2007.

[29] Yong Jiang and Jochen C. Reif. Efficient algorithms for calculating epistatic genomic relationship matrices. Genetics, 216(3):651–669, 2020.

