## Supplementary Data File for "Fast and Accurate Exhaustive Higher-Order Epistasis Search with BitEpi"

Arash Bayat*, Brendan Hosking, Yatish Jain
Cameron Hosking, Milindi Kodikara, Daniel Reti,
Natalie Twine and Denis C. Bauer*

**Health and Biosecurity, CSIRO, Australia**

*
*

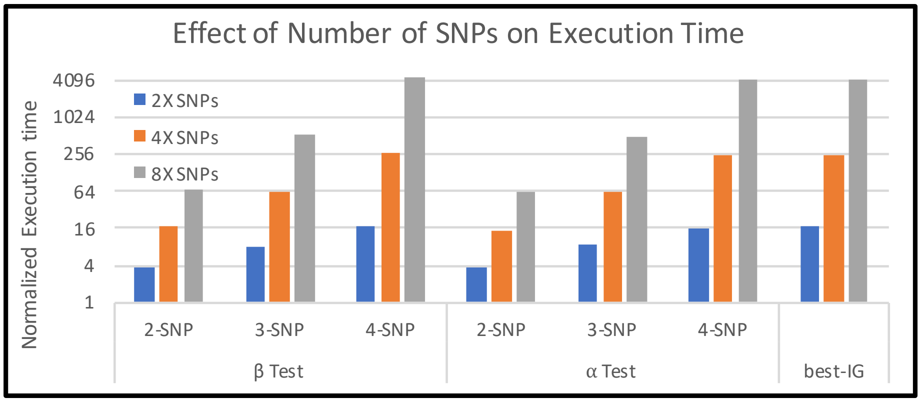


(a) The effect of number of SNPs on BitEpi execution time.

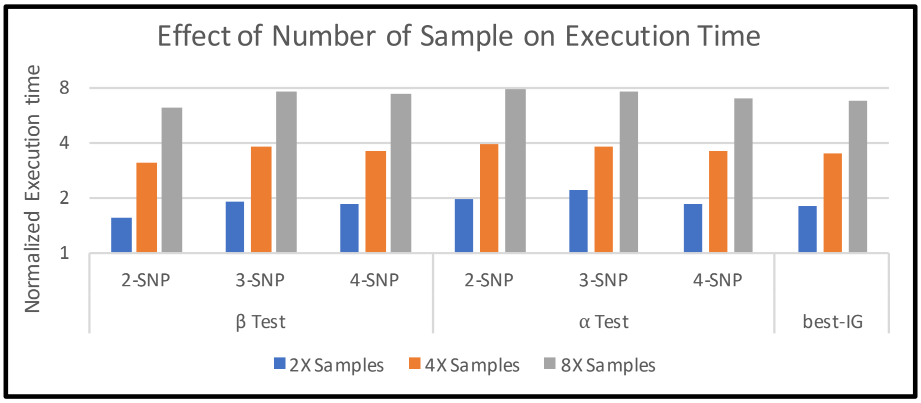


(b) The effect of number of samples on BitEpi execution time.

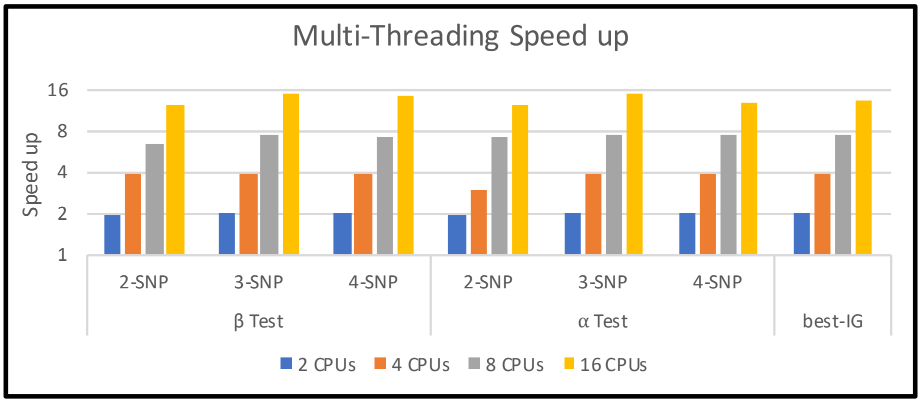


(c) The effect of number of threads on BitEpi execution time.

Figure 1: The effect of number of SNPs, samples and threads on BitEpi execution time.

Table 1: BitEpi execution time when the number of SNPs varies in the dataset. The number of samples is 4,000 (2,000 cases and 2,000 controls) in all cases.

| Test | Number of SNP | Execution Time (sec) |
|---|---:|---:|
| best test | 50 | 0.82 |
| | 100 | 13.65 |
| | 200 | 207.58 |
| | 400 | 3320.27 |
| $\alpha$ Test 2-SNP | 5000 | 40.19 |
| | 10000 | 155.33 |
| | 20000 | 611.36 |
| | 40000 | 2423.67 |
| $\alpha$ Test 3-SNP | 250 | 8.04 |
| | 500 | 72.97 |
| | 1000 | 506.50 |
| | 2000 | 4047.00 |
| $\alpha$ Test 4-SNP | 50 | 0.82 |
| | 100 | 12.91 |
| | 200 | 207.16 |
| | 400 | 3327.03 |
| $\beta$ Test 2-SNP | 5000 | 40.19 |
| | 10000 | 155.27 |
| | 20000 | 688.16 |
| | 40000 | 2711.73 |
| $\beta$ Test 3-SNP | 250 | 7.94 |
| | 500 | 63.11 |
| | 1000 | 504.49 |
| | 2000 | 4034.00 |
| $\beta$ Test 4-SNP | 50 | 0.75 |
| | 100 | 12.59 |
| | 200 | 202.91 |
| | 400 | 3281.52 |

Table 2: BitEpi execution time when the number of samples varies in the dataset.

| Test | Number of SNP | Number of Sample | Execution Time (sec) |
|---|---|---|---|
| best test | 200 | 2000 | 115.90 |
| | 200 | 4000 | 207.77 |
| | 200 | 8000 | 400.76 |
| | 200 | 16000 | 785.50 |
| $\alpha$ Test 2-SNP | 10000 | 2000 | 79.07 |
| | 10000 | 4000 | 155.62 |
| | 10000 | 8000 | 310.64 |
| | 10000 | 16000 | 618.94 |
| $\alpha$ Test 3-SNP | 500 | 2000 | 33.07 |
| | 500 | 4000 | 73.08 |
| | 500 | 8000 | 125.68 |
| | 500 | 16000 | 249.03 |
| $\alpha$ Test 4-SNP | 200 | 2000 | 111.88 |
| | 200 | 4000 | 207.26 |
| | 200 | 8000 | 408.13 |
| | 200 | 16000 | 785.06 |
| $\beta$ Test 2-SNP | 10000 | 2000 | 100.42 |
| | 10000 | 4000 | 155.53 |
| | 10000 | 8000 | 310.31 |
| | 10000 | 16000 | 618.24 |
| $\beta$ Test 3-SNP | 500 | 2000 | 32.83 |
| | 500 | 4000 | 63.09 |
| | 500 | 8000 | 124.77 |
| | 500 | 16000 | 247.52 |
| $\beta$ Test 4-SNP | 200 | 2000 | 109.28 |
| | 200 | 4000 | 203.11 |
| | 200 | 8000 | 392.05 |
| | 200 | 16000 | 814.35 |

Table 3: BitEpi execution time when the number of threads varies. The number of samples is 4,000 (2,000 cases and 2,000 controls) in all cases.

| Test | Number of SNP | Number of Threads | Execution Time (sec) |
|---|---|---|---|
| best Test | 400 | 1 | 3158.39 |
| | 400 | 2 | 1589.64 |
| | 400 | 4 | 811.48 |
| | 400 | 8 | 425.12 |
| | 400 | 16 | 236.35 |
| $\alpha$ Test 2-SNP | 40000 | 1 | 2370.38 |
| | 40000 | 2 | 1200.37 |
| | 40000 | 4 | 781.90 |
| | 40000 | 8 | 323.10 |
| | 40000 | 16 | 188.89 |
| $\alpha$ Test 3-SNP | 2000 | 1 | 3844.00 |
| | 2000 | 2 | 1927.50 |
| | 2000 | 4 | 973.27 |
| | 2000 | 8 | 503.19 |
| | 2000 | 16 | 255.58 |
| $\alpha$ Test 4-SNP | 400 | 1 | 3154.43 |
| | 400 | 2 | 1586.92 |
| | 400 | 4 | 806.27 |
| | 400 | 8 | 423.24 |
| | 400 | 16 | 244.00 |
| $\beta$ Test 2-SNP | 40000 | 1 | 2653.23 |
| | 40000 | 2 | 1341.05 |
| | 40000 | 4 | 685.57 |
| | 40000 | 8 | 405.74 |
| | 40000 | 16 | 214.14 |
| $\beta$ Test 3-SNP | 2000 | 1 | 3840.00 |
| | 2000 | 2 | 1917.02 |
| | 2000 | 4 | 970.26 |
| | 2000 | 8 | 504.73 |
| | 2000 | 16 | 255.69 |
| $\beta$ Test 4-SNP | 400 | 1 | 3125.74 |
| | 400 | 2 | 1571.14 |
| | 400 | 4 | 801.53 |
| | 400 | 8 | 425.05 |
| | 400 | 16 | 219.61 |

Table 4: Description of epistasis model-simulated with GAMETES and 2-SNP detection power of BOOST and BitEpi for each model. For each model, 100 datasets are generated each with 100 SNPs and 2,000 samples (1,000 case and 1,000 controls). The minor allele frequency of SNPs in each dataset varies between 0.01 and 0.5. MAF1 and MAF2 are the minor allele frequency of the first and second interactive SNPs. Detection power is the number of datasets (out of 100) where the interactive pair is ranked first by the analysis metod.

| | Model Characteristics | | | Detection Power | |
|---|---|---|---|---|---|
| Model | Heritability | MAF1 | MAF2 | BOOST | BitEpi |
| PM1 | 0.005 | 0.050 | 0.050 | 10 | 6 |
| PM2 | 0.005 | 0.050 | 0.250 | 66 | 81 |
| PM3 | 0.005 | 0.050 | 0.500 | 45 | 55 |
| PM4 | 0.005 | 0.250 | 0.250 | 68 | 88 |
| PM5 | 0.005 | 0.250 | 0.500 | 55 | 79 |
| PM6 | 0.005 | 0.500 | 0.500 | 64 | 82 |
| PM7 | 0.050 | 0.050 | 0.050 | 70 | 100 |
| PM8 | 0.050 | 0.050 | 0.250 | 75 | 100 |
| PM9 | 0.050 | 0.250 | 0.250 | 71 | 100 |
| PM10 | 0.200 | 0.500 | 0.500 | 75 | 100 |

Table 5: Description of epistasis model-simulated with GAMETES and 3-SNP detection power of MPI3SNP and BitEpi for each model. For each model, 100 datasets are generated each with 100 SNPs and 2,000 samples (1,000 cases and 1,000 controls). The minor allele frequency of SNPs in each dataset varies between 0.01 and 0.5. MAF1, MAF2, and MAF3 are the minor allele frequency of the first, second and third interactive SNPs. Detection power is the number of datasets (out of 100) where the interactive 3-SNP is ranked first by the analysis method.

| | Model Characteristics | | | | Detection Power | |
|---|---|---|---|---|---|---|
| Model | Heritability | MAF 1 | MAF 2 | MAF 3 | MPI3SNP | BitEpi |
| TM1 | 0.005 | 0.050 | 0.050 | 0.050 | 0 | 0 |
| TM2 | 0.005 | 0.050 | 0.250 | 0.500 | 16 | 25 |
| TM3 | 0.005 | 0.050 | 0.500 | 0.500 | 9 | 16 |
| TM4 | 0.005 | 0.250 | 0.250 | 0.250 | 13 | 36 |
| TM5 | 0.005 | 0.250 | 0.250 | 0.500 | 15 | 36 |
| TM6 | 0.005 | 0.500 | 0.500 | 0.500 | 27 | 52 |
| TM7 | 0.050 | 0.250 | 0.250 | 0.250 | 100 | 100 |
| TM8 | 0.050 | 0.250 | 0.250 | 0.500 | 100 | 100 |
| TM9 | 0.050 | 0.500 | 0.500 | 0.500 | 100 | 100 |

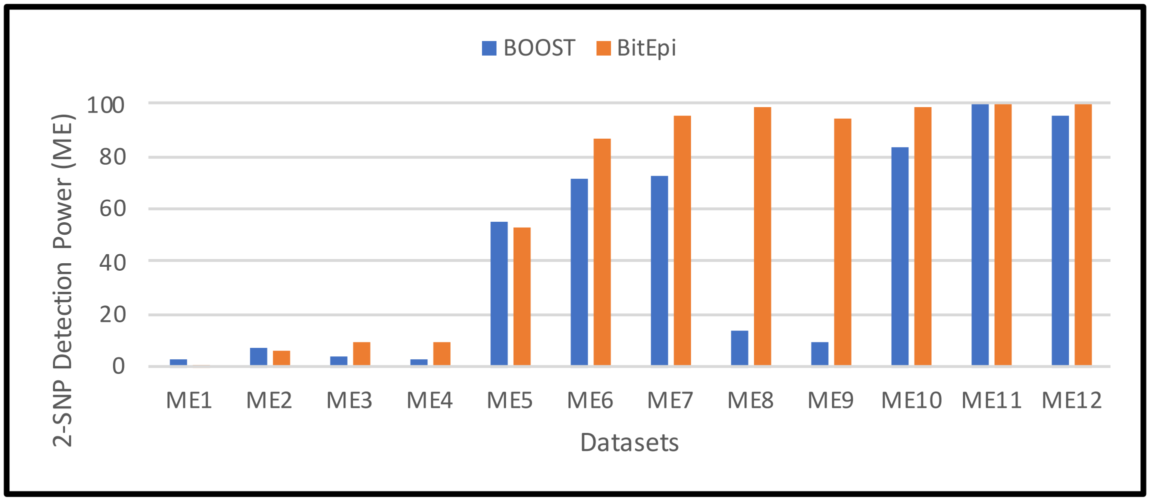


(a) The effect of number of SNPs on BitEpi execution time.

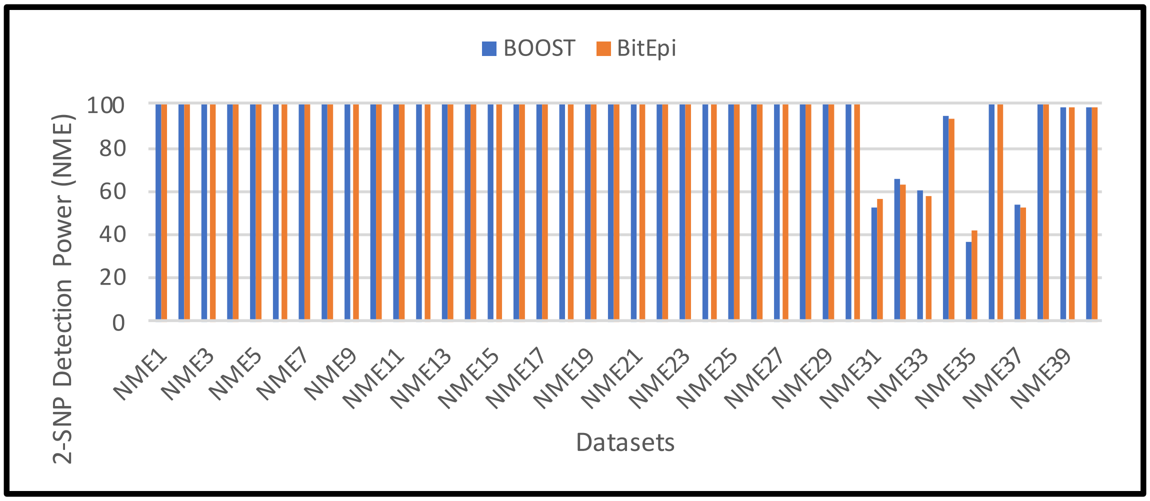


(b) The effect of number of samples on BitEpi execution time.

Figure 2: The effect of number of SNPs, samples and threads on BitEpi execution time.

Table 6: Description of ME epistasis models simulated with GAMETES in [1] and 2-SNP detection power of BOOST and BitEpi for each model. For each model, 100 datasets are generated each with 100 SNPs and 1,600 samples (800 cases and 800 controls). Detection power is the number of datasets (out of 100) where the interactive pair is ranked first by the analysis method.

| Model Identifier (Downloaded Data) | Model | Detection Power | |
|---|---|---|---|
| | | BOOST | BitEpi |
| 70 | ME1 | 3 | 1 |
| 71 | ME2 | 7 | 6 |
| 72 | ME3 | 4 | 9 |
| 73 | ME4 | 3 | 9 |
| 74 | ME5 | 55 | 53 |
| 75 | ME6 | 72 | 87 |
| 76 | ME7 | 73 | 96 |
| 77 | ME8 | 14 | 99 |
| 78 | ME9 | 10 | 94 |
| 79 | ME10 | 84 | 99 |
| 80 | ME11 | 100 | 100 |
| 81 | ME12 | 96 | 100 |

Table 7: Description of NME epistasis models simulated with GAMETES in and 2-SNP detection power of BOOST and BitEpi for each model. For each model, 100 datasets are generated each with 100 SNPs and 1,600 samples (800 cases and 800 controls). Detection power is the number of datasets (out of 100) where the interactive pair is ranked first by the analysis method.

| Model Identifier (Downloaded Data) | Model | Detection Power | |
|---|---|---|---|
| | | BOOST | BitEpi |
| 0 | NME1 | 100 | 100 |
| 1 | NME2 | 100 | 100 |
| 2 | NME3 | 100 | 100 |
| 3 | NME4 | 100 | 100 |
| 4 | NME5 | 100 | 100 |
| 5 | NME6 | 100 | 100 |
| 6 | NME7 | 100 | 100 |
| 7 | NME8 | 100 | 100 |
| 8 | NME9 | 100 | 100 |
| 9 | NME10 | 100 | 100 |
| 15 | NME11 | 100 | 100 |
| 16 | NME12 | 100 | 100 |
| 17 | NME13 | 100 | 100 |
| 18 | NME14 | 100 | 100 |
| 19 | NME15 | 100 | 100 |
| 25 | NME16 | 100 | 100 |
| 26 | NME17 | 100 | 100 |
| 27 | NME18 | 100 | 100 |
| 28 | NME19 | 100 | 100 |
| 29 | NME20 | 100 | 100 |

| Model Identifier (Downloaded Data) | Model | Detection Power | |
|---|---|---|---|
| | | BOOST | BitEpi |
| 30 | NME21 | 100 | 100 |
| 31 | NME22 | 100 | 100 |
| 32 | NME23 | 100 | 100 |
| 33 | NME24 | 100 | 100 |
| 34 | NME25 | 100 | 100 |
| 40 | NME26 | 100 | 100 |
| 41 | NME27 | 100 | 100 |
| 42 | NME28 | 100 | 100 |
| 43 | NME29 | 100 | 100 |
| 44 | NME30 | 100 | 100 |
| 55 | NME31 | 52 | 57 |
| 56 | NME32 | 66 | 63 |
| 57 | NME33 | 60 | 58 |
| 58 | NME34 | 95 | 93 |
| 59 | NME35 | 37 | 42 |
| 65 | NME36 | 100 | 100 |
| 66 | NME37 | 54 | 53 |
| 67 | NME38 | 100 | 100 |
| 68 | NME39 | 99 | 99 |
| 69 | NME40 | 99 | 99 |

Table 8: Overview of the tools used in this study

| Tool | max. interactions |
|---|---|
| Boost | 2 |
| MPI3SNP | 3 |
| MDR | 4 |
| BitEpi | 4 |

### Proof of increasing $\beta$

Here we would like to prove that when adding a SNP to an interaction the $\beta$ value always either increases or remains the same. For example, interaction ABC has an equal or higher $\beta$ than interaction AB ($\beta_{ABC} \geq \beta_{AB}$) where A, B and C are three SNPs. As described in the manuscript, there are $3^m$ rows in the contingency table of an $m$-SNP interaction. Therefore in an $(m+1)$-SNP interaction there are $3^{(m+1)}$ rows ($3 \times 3^m$ rows). In other words, each row of the $m$-SNP interaction is divided into 3 rows in the $(m+1)$-SNP interaction. Table 9 clarifies this mapping, where $x = \sum_{i=1}^{3} x_i$ and $y = \sum_{i=1}^{3} y_i$ (i.e. the numbers of cases and controls with A=0/1 and B=1/1 do not change when adding C to the interaction).

Table 9: For $m = 2$: an $m$-SNP interaction on the left, an $(m+1)$-SNP interaction on the right, and the row mapping in the middle. The table shows that the "A=0/1, B=1/1" row is divided into 3 rows when adding C to the interaction. Note that the total number of cases and controls are the same.

| $m$-SNP interaction | | | | | a $m$-SNP row $\longrightarrow$ 3 $(m+1)$-SNP row | | | | | $(m+1)$-SNP interaction | | | | |
|---|---|---|---|---|---|---|---|---|---|---|---|---|---|---|
| A | B | Case | Control | | AB | C | Cases | Controls | | A | B | C | Cases | Controls |
| 0/0 | 0/0 | ... | ... | | 0/0 . 0/0 | 0/0 | ... | ... | | 0/0 | 0/0 | 0/0 | ... | ... |
| 0/0 | 0/1 | ... | ... | | ... | ... | ... | ... | | ... | ... | ... | ... | ... |
| ... | ... | ... | ... | | | **0/0** | $x_1$ | $y_1$ | | **0/1** | **1/1** | **0/0** | $x_1$ | $y_1$ |
| **0/1** | **1/1** | $x$ | $y$ | | **0/1 . 1/1** | **0/1** | $x_2$ | $y_2$ | | **0/1** | **1/1** | **0/1** | $x_2$ | $y_2$ |
| ... | ... | ... | ... | | | **1/1** | $x_3$ | $y_3$ | | **0/1** | **1/1** | **1/1** | $x_3$ | $y_3$ |
| 1/1 | 0/1 | ... | ... | | ... | ... | ... | ... | | ... | ... | ... | ... | ... |
| 1/1 | 1/1 | ... | v | | 1/1 . 1/1 | 1/1 | ... | ... | | 1/1 | 1/1 | 1/1 | ... | ... |

$\beta$ is the weighted average of the purity of all rows. Let $K_r$ be the $r^{th}$ row of the $m$-SNP interaction with $x$ cases and $y$ controls. Let $K3_r$ be the corresponding 3 rows in the $(m+1)$-SNP interaction, where $x_i$ and $y_i$ represent the number of cases and controls in the $i^{th}$ row (i.e. $x_1, x_2, x_3, y_1, y_2, y_3$). To show $\beta_{ABC} \geq \beta_{AB}$ it is enough to show that for each row in an $m$-SNP interaction ($r$) the contribution of $K3_r$ to $(m+1)$-SNP $\beta$ (i.e. $\beta_{ABC}$ in Table 9) is higher than the contribution of $K_r$ to $m$-SNP $\beta$ (i.e. $\beta_{AB}$ in Table 9). We represent the former contribution with $P_{m+1}$ (Equation 4) and the latter contribution with $P_m$ (Equation 3). $S$ is the total number of samples. Therfore we should prove Inequality 5. We expand and simplify this inequality in several steps (shown below) to form Inequality 13. It can be seen that Inequality 13 holds true as $x_i \geq 0$ and $y_i \geq 0$ and the possible negative values are squared.

$$x = \sum_{i=1}^{3} x_i \tag{1}$$

$$y = \sum_{i=1}^{3} y_i \tag{2}$$

$$P_m = \frac{x+y}{S} \times \frac{x^2+y^2}{(x+y)^2} \\ = \frac{1}{S} \times \frac{x^2+y^2}{(x+y)} \tag{3}$$

$$P_{m+1} = \sum_{i=1}^{3} (\frac{x_i+y_i}{S}) \times (\frac{{x_i}^2+{y_i}^2}{(x_i+y_i)^2}) \\ = \frac{1}{S} \times \sum_{i=1}^{3} \frac{{x_i}^2+{y_i}^2}{(x_i+y_i)} \tag{4}$$

$$P_{m+1} \geq P_m \tag{5}$$

$$\frac{1}{S} \times \sum_{i=1}^{3} \frac{{x_i}^2+{y_i}^2}{(x_i+y_i)} \geq \frac{1}{S} \times \frac{x^2+y^2}{(x+y)} \tag{6}$$

$$\sum_{i=1}^{3} \frac{{x_i}^2+{y_i}^2}{(x_i+y_i)} \geq \frac{x^2+y^2}{x+y} \tag{7}$$

$$\sum_{i=1}^{3} \frac{{x_i}^2+{y_i}^2}{(x_i+y_i)} \geq \frac{(\sum_{i=1}^{3} x_i)^2 + (\sum_{i=1}^{3} y_i)^2}{\sum_{i=1}^{3} x_i + \sum_{i=1}^{3} y_i} \tag{8}$$

$$\frac{{x_1}^2+{y_1}^2}{(x_1+y_1)} + \frac{{x_2}^2+{y_2}^2}{(x_2+y_2)} + \frac{{x_3}^2+{y_3}^2}{(x_3+y_3)} \geq \frac{(x_1+x_2+x_3)^2+(y_1+y_2+y_3)^2}{x_1+x_2+x_3+y_1+y_2+y_3} \tag{9}$$

$$x_1x_2^2y_3^2 + x_1y_2^2x_3^2 + y_1x_2^2y_3^2 + y_1y_2^2x_3^2 + x_1^2x_2y_3^2 + y_1^2x_2x_3^2 + \\ x_1^2y_2y_3^2 + y_1^2y_2x_3^2 + x_1^2y_2^2x_3 + y_1^2x_2^2x_3 + x_1^2y_2^2y_3 + y_1^2x_2^2y_3 \geq \\ 2x_1x_2x_3y_1y_2 + 2x_1x_2x_3y_1y_3 + 2x_1x_2x_3y_2y_3 + \\ 2x_1x_2y_1y_2y_3 + 2x_1x_3y_1y_2y_3 + 2x_2x_3y_1y_2y_3 \tag{10}$$

$$x_1x_2^2y_3^2 + x_1y_2^2x_3^2 + y_1x_2^2y_3^2 + y_1y_2^2x_3^2 + \\ x_1^2x_2y_3^2 + y_1^2x_2x_3^2 + x_1^2y_2y_3^2 + y_1^2y_2x_3^2 + \\ x_1^2y_2^2x_3 + y_1^2x_2^2x_3 + x_1^2y_2^2y_3 + y_1^2x_2^2y_3 - \\ 2x_1x_2x_3y_1y_2 - 2x_1x_2x_3y_1y_3 - 2x_1x_2x_3y_2y_3 - \\ 2x_1x_2y_1y_2y_3 - 2x_1x_3y_1y_2y_3 - 2x_2x_3y_1y_2y_3 \geq 0 \tag{11}$$

$$
\begin{aligned}
(x_1x_2^2y_3^2 - 2x_1x_2x_3y_2y_3 + x_1y_2^2x_3^2)+ \\
(y_1x_2^2y_3^2 - 2x_2x_3y_1y_2y_3 + y_1y_2^2x_3^2)+ \\
(x_1^2x_2y_3^2 - 2x_1x_2x_3y_1y_3 + y_1^2x_2x_3^2)+ \\
(x_1^2y_2y_3^2 - 2x_1x_3y_1y_2y_3 + y_1^2y_2x_3^2)+ \\
(x_1^2y_2^2x_3 - 2x_1x_2x_3y_1y_2 + y_1^2x_2^2x_3)+ \\
(x_1^2y_2^2y_3 - 2x_1x_2y_1y_2y_3 + y_1^2x_2^2y_3) \geq 0
\end{aligned} \tag{12}
$$

$$
\begin{aligned}
x_1(x_2y_3 - y_2x_3)^2+ \\
y_1(x_2y_3 - y_2x_3)^2+ \\
x_2(x_1y_3 - y_1x_3)^2+ \\
y_2(x_1y_3 - y_1x_3)^2+ \\
x_3(x_1y_2 - y_1x_2)^2+ \\
y_3(x_1y_2 - y_1x_2)^2 \geq 0
\end{aligned} \tag{13}
$$

### References

[1] Peng-Jie Jing and Hong-Bin Shen. Macoed: a multi-objective ant colony optimization algorithm for snp epistasis detection in genome-wide association studies. *Bioinformatics*, 31(5):634–641, 2014.
